# An integrated approach to identify environmental modulators of genetic risk factors for complex traits

**DOI:** 10.1101/2021.02.23.432608

**Authors:** Brunilda Balliu, Ivan Carcamo -Orive, Michael J. Gloudemans, Daniel C. Nachun, Matthew G. Durrant, Steven Gazal, Chong Y. Park, David A. Knowles, Martin Wabitsch, Thomas Quertermous, Joshua W. Knowles, Stephen B. Montgomery

## Abstract

Complex traits and diseases can be influenced by both genetics and environment. However, given the large number of environmental stimuli and power challenges for gene-by-environment testing, it remains a critical challenge to identify and prioritize specific disease-relevant environmental exposures. We propose a novel framework for leveraging signals from transcriptional responses to environmental perturbations to identify disease-relevant perturbations that can modulate genetic risk for complex traits and inform the functions of genetic variants associated with complex traits. We perturbed human skeletal muscle, fat, and liver relevant cell lines with 21 perturbations affecting insulin resistance, glucose homeostasis, and metabolic regulation in humans and identified thousands of environmentally responsive genes. By combining these data with GWAS from 31 distinct polygenic traits, we show that heritability of multiple traits is enriched in regions surrounding genes responsive to specific perturbations and, further, that environmentally responsive genes are enriched for associations with specific diseases and phenotypes from the GWAS catalogue. Overall, we demonstrate the advantages of large-scale characterization of transcriptional changes in diversely stimulated and pathologically relevant cells to identify disease-relevant perturbations.

## Introduction

Genome-wide association studies (GWAS) have identified thousands of genetic variants associated with complex diseases and traits^1^. The majority of these variants fall into non-coding regions of the genome and, as a result, their mechanism of action remains largely unknown^2^. In recent years, researchers have gained an increasingly clear picture of which parts of the genome are active in a range of tissues and cell types^3–6^. Integrating such information with results from GWAS has identified cell types, tissues, and regulatory elements relevant to specific diseases and phenotypes and moved the field towards mechanistic understanding of GWAS hits^7–9^. In addition, genomic colocalization and transcriptome-wide association studies combining results from GWAS and expression quantitative trait loci (eQTL) studies have identified candidate causal genes and their mechanisms of action^10–12^.

Despite these advances, a modest fraction of GWAS associated variants and eQTLs colocalize for any trait^13,14^ providing the perspective that many disease-relevant effects are modulated by yet-to-be-discovered environmental factors. To address this challenge, multiple studies have mapped eQTLs in vitro that are responsive to the environment^15–26^. For example, the Immune Variation project identified eQTLs in human CD4+ T lymphocytes with different effects across distinct immune states^17^. These previously unknown, immune state-specific eQTLs were enriched for autoimmune disease-associated variants, underscoring the importance of exploring contexts beyond tissues and cell types to reveal the specificity of genetic associations. Although there is mounting evidence that environment modulates genetic effects, GWAS and eQTL studies rarely measure and test for genetic interactions with environment exposures. This is, in part, due to the difficulty of identifying and collecting information on the most relevant environmental exposures in GWAS cohorts and performing eQTL studies in contexts that are relevant for the specific trait or disease.

In this study, we extend the current understanding of inherited variation in complex traits by implementing a novel framework to model signals from transcriptional responses to environmental perturbations in order to identify and prioritize disease-relevant environments that can modulate genetic risk for complex traits and inform the functions of genetic variants and genes associated with complex traits. Specifically, we first assessed environmental effects on gene expression levels in three metabolic human cell lines by performing RNA-seq in muscle-, fat-, and liver-relevant cell lines treated with 21 different environmental perturbations related to aspects of glucose and insulin metabolism, kinase inhibitors, inflammation, fatty acid metabolism, etc. (N=234 samples). We identified thousands of environmentally responsive genes underlying disease-associated response pathways and characterized the specificity and sharing of these effects across perturbations and cell lines. Next, to identify disease-relevant perturbations, we coupled our gene expression data with GWAS summary statistics of 31 complex traits and diseases as well as associations from the GWAS catalogue. We confirmed several well-established environmental-phenotype associations, e.g., the role of TGF-β1 on asthma^27^ and provided additional evidence for recent and less well-understood associations, e.g., the role of leptin on major depressive disorder^28^. Last, to further illustrate how perturbation experiments inform the functions of complex trait associated variants, we integrate our perturbation data with genomic colocalization studies and show that the effects of these perturbations in the relevant tissues identifies context-specific molecular mechanisms of GWAS hits for diverse cardiometabolic traits.

This resource characterizes the dynamic transcriptional landscape in metabolic tissues and provides a framework to identify and prioritize disease-relevant perturbations and disentangle the complex gene-environment interactions that determine disease susceptibility, which is particularly relevant for complex traits such as insulin resistance, diabetes and obesity.

## Results

### Transcriptome map of 21 perturbations across human skeletal muscle, fat and liver cell lines

We generated a transcriptome map of multiple chemical and environmental perturbations in well-established human skeletal muscle, fat and liver cell lines (N=234 samples). Specifically, we studied 21 environmental perturbations covering multiple aspects of glucose and insulin metabolism, inflammation, fatty acid metabolism, and including both LDL-lowering and anti-diabetic drugs (Figure 1 and Table S1). For each perturbation and cell line and matched controls, we conducted assays in triplicate and applied differential expression analysis. We observed that the majority of perturbations induced broad gene expression changes in at least one cell line at FDR < 5% (Figure 1A, Table S2). Several perturbations induced broad changes across all cell lines; for example, insulin and IGF1 altered the gene expression of 1,500-2,000 genes in each cell line. Other perturbations had broad changes only in specific cell lines. For example, IL-6, lauroyl-l-carnitine, and glucose had more pronounced effects in fat, muscle, and liver, respectively, impacting the expression of 3,161, 2,051 and 2,724 genes, respectively.

**Figure 1.**
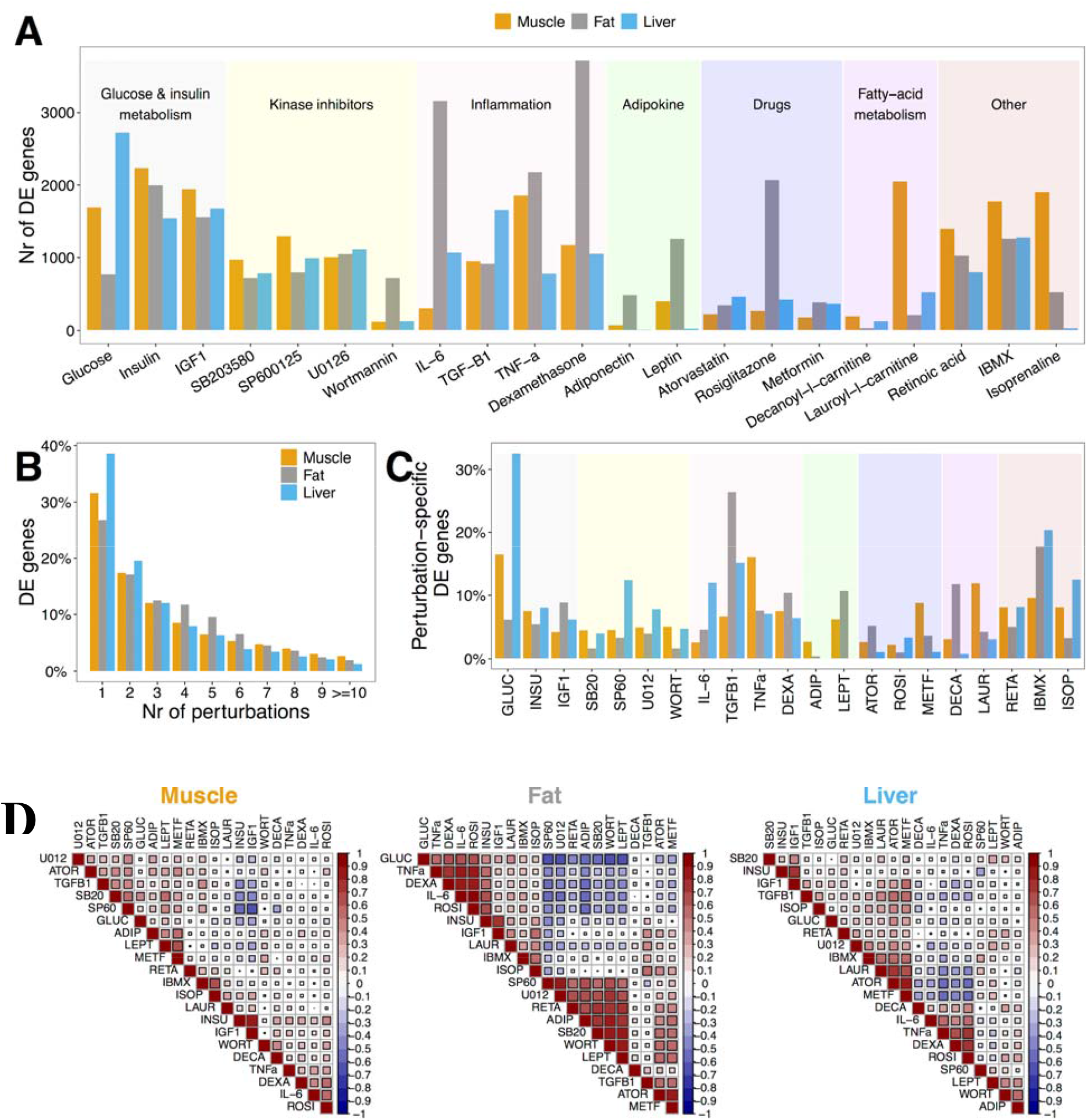
Perturbations induce large-scale changes in gene expression in muscle, fat, and liver. **(A)** Number of DE genes for each perturbation in each cell line (FDR<5%). See Table S2 for extended DE summary statistics. **(B)** Proportion of DE genes that change in response to up to 10 perturbations in each cell line. See Tables S4 for extended results on sharing of DE genes. **(C)** Proportion of perturbation-specific DE genes, i.e., genes that change in response to a single perturbation, within each cell line. **(D)** Correlation of DE patterns between different perturbations within each cell line. Each square is Spearman’s correlation between the DE test statistic of a pair of perturbations across all genes. DE: differentially expressed; FDR: False Discovery Rate.

Despite the broad effects for each perturbation, multiple differentially expressed (DE) genes showed perturbation-specific effects within each cell line, highlighting a unique molecular response to each perturbation. We observed 1,883 genes in muscle, 1,813 genes in fat, and 2,231 genes in liver altered by only a single perturbation in their respective cell lines (Figure 1B and Table S3). The largest proportions of perturbation-specific DE genes were found in glucose-stimulated liver cell lines and TGF-β1-stimulated fat cell lines. For these perturbations, 32.6% and 26.4% of DE genes were not altered by any of the other 20 perturbations in the same cell line (Figure 1C). By further stratifying across these cell lines, we identified 627, 742, and 808 genes that were both perturbation- and cell line-specific DE genes in muscle, fat, and liver (FDR < 5%; Figure S4A and Table S3). Glucose-stimulated liver cells also provided the largest amount of perturbation-and cell line-specific DE genes; 9.8% of DE genes were not altered by any of the other 20 perturbations in any cell line or by glucose stimulation in fat or muscle.

To identify the relationships between perturbations based on their overall transcriptional responses, we assessed the correlation of DE genes between each pair of perturbations within the same cell line (Figures 1D). The correlation of the effect of some perturbations was similar across cell lines, e.g., the effects of insulin and IGF1 were positively correlated in all three cell lines, i.e., Spearman’s ρ = 0.88, 0.76, and 0.71 in muscle, fat, and liver, respectively. The relationship of other perturbations, however, was dependent on the cellular context, e.g., while the effects of glucose and wortmannin were moderately correlated in fat (Spearman’s ρ = -0.63), their correlation in muscle and liver was low (Spearman’s ρ =0.02 and 0.2, respectively).

To explore the shared and specific pathways altered by each perturbation, we performed enrichment analysis of DE genes in annotated pathways from ConsensusPathDB^29^ (Table S3). Our analysis highlighted multiple shared pathways across perturbations and cell lines related to PI3K-AKT-mTOR, MAPK, adipogenesis, and TGF-β signaling (Figure S4B). We also observed several differences in pathway enrichments; for example, pathways related to FOXA2 and FOXA3 transcription factor networks had greater enrichment across several perturbations in liver than in muscle and fat, transcriptional regulation by RUNX2 had greater enrichment in muscle than in liver and fat, and chromatin organization and remodeling pathways had greater enrichment in fat than in liver and muscle. In addition, for genes affected by multiple perturbations we saw strong enrichment pathways related to insulin signaling and resistance.

Combined, our concurrent assessment of multiple metabolically relevant perturbations across cell lines highlights the relationships between complex cell-specific molecular mechanisms and provides a genome-wide map of genes and signaling pathways with potential environmental contributions to complex disease susceptibility.

### Prioritizing complex disease-relevant environmental perturbations

To measure the relevance of diverse environmental perturbations in complex diseases, we analyzed our transcriptome data together with GWAS summary statistics for 31 diseases and complex traits broadly related to multiple cardiometabolic, psychiatric, autoimmune, and reproductive traits, as well as hematological measurements (Figure 2 and S5; Table S5). We hypothesized that environmental perturbations impact disease through the same genes that confer susceptibility to the trait. To this end, for each of the 21 perturbations across the three cell lines, we used stratified LD score regression^8,9,30^ (LDSCreg) to test whether disease heritability, i.e., proportion of phenotypic variance determined by genotypic variance, is enriched in regions surrounding DE genes for that perturbation and cell line, adjusting for both heritability explained by a baseline model of genetic architecture^9^ and by regions surrounding genes expressed in the specific cell line.

**Figure 2.**
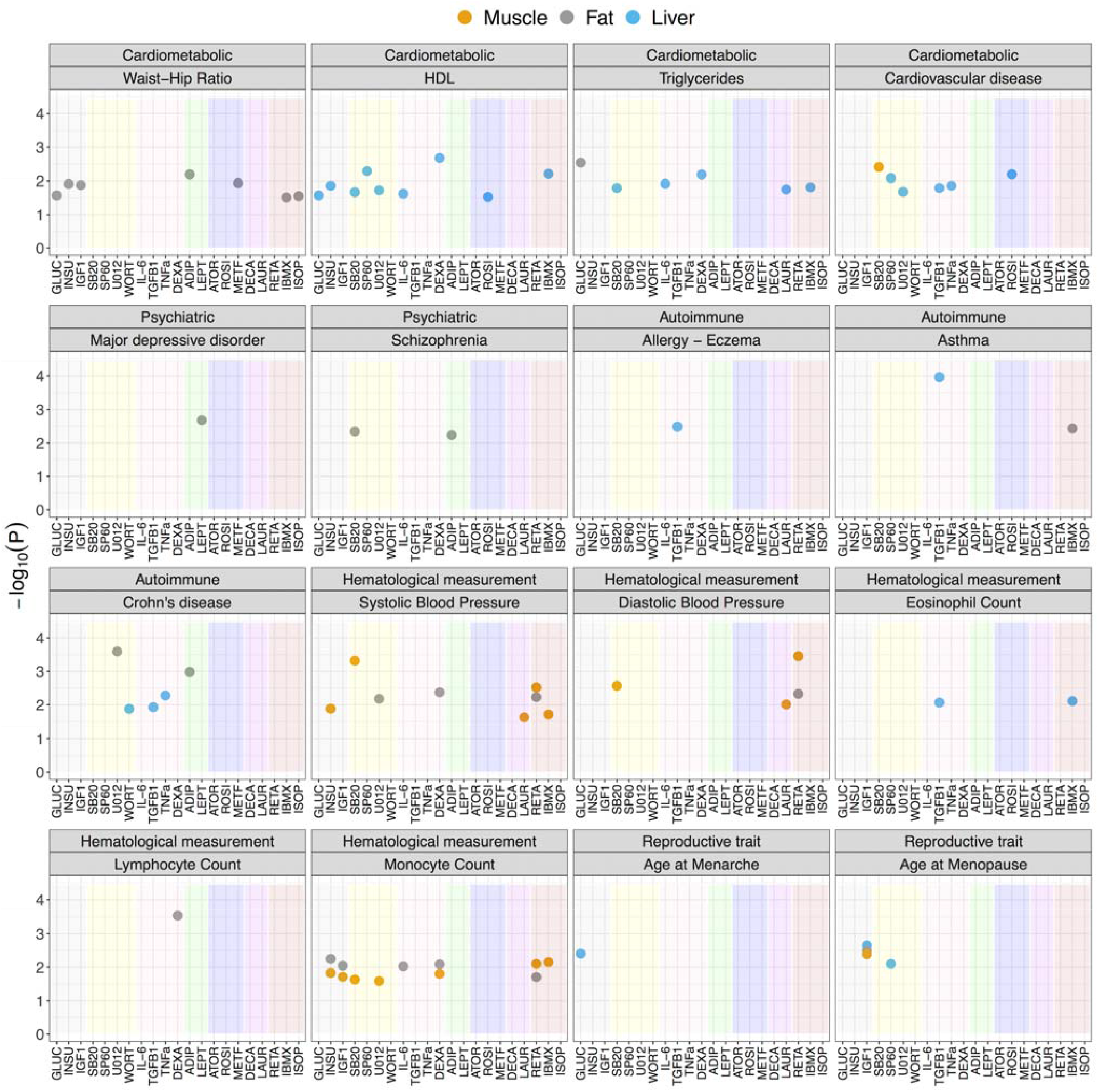
Prioritizing complex disease-relevant environmental perturbations via heritability enrichment analysis. Heritability enrichment results for each complex trait. Each point represents a perturbation-cell-line combination that passes the FDR<10% cut off. The y-axis represents the -log_10_(P-value) of heritability enrichment, the x-axis indicates perturbation, color of the point indicates cell line, and the shading color within each panel indicates the perturbation category from Figure 1A. Numerical results are reported in Table S5.

For 26 of the 31 traits tested, the SNP-based heritability estimate was sufficiently large to partition reliably with LDSCreg, i.e., heritability Z-score >= 7 (Table S5). In 19 of these traits, at least one perturbation in at least one cell line was enriched for heritability (FDR<10%; Figure 2). Several of the enrichments recapitulate important known biology. For example, among cardiometabolic traits, high-density lipoprotein (HDL) and triglyceride levels were enriched for dexamethasone (P=2.10 × 10^−3^ and P=6.53 × 10^−3^), a corticosteroid known to induce dyslipidemia^31,32^, and cardiovascular disease was enriched for rosiglitazone (P=6.44 × 10^−3^), an antidiabetic drug shown to increase risk of cardiovascular disease^33^. In addition, these enrichments were often manifested through a single specific relevant cell line. For example, waist-hip ratio (WHR) heritability was enriched for genes whose expression is modified by perturbations in fat, while triglyceride and HDL level heritability were enriched for genes whose expression is modified by perturbations in the liver.

Several notable examples were also observed for other tested traits. For psychiatric disorders, leptin, a hormone produced and secreted by white adipose tissue that is associated with antidepressant-like actions^28,34,35^, was enriched for heritability of major depressive disorder via its effect in fat cell lines (P=2.11 × 10^−3^). In addition, adiponectin, plasma levels of which appear to be altered in neurological disorders with metabolic and inflammatory components^36–38^, was enriched for heritability of schizophrenia (P=5.81 × 10^−3^). For tested autoimmune diseases, TGF-β1, an immune-suppressive cytokine dysregulated in the intestines of inflammatory bowel disease patients^39^, was enriched for heritability of Crohn’s disease (P=1.18 × 10^−2^), as well as heritability of allergy, eczema, and asthma^27,40–42^ (P=1.08 × 10^−4^), three diseases with shared genetic origin^43^. Several perturbations were also enriched for heritability of hematological measurements; for example, dexamethasone, a synthetic glucocorticoid known to deplete peripheral blood lymphocytes and impact immune response^44^, was enriched for heritability of lymphocyte count (P=2.88 × 10^−4^). Lastly, for reproductive traits, glucose was enriched for heritability of age at menarche -older age at menarche is associated with reduced risk of glucose metabolism disorder^45^ -while IGF1, whose serum levels rapidly decrease after menopause^46^, was enriched for heritability of age at menopause (P=2.21 × 10^−3^).

### Identifying environmental perturbations impacting GWAS-significant loci

Beyond the broad polygenic impact of the tested perturbations and in order to analyze a larger number of traits, we sought to prioritize the subset of perturbations that were enriched for impact on GWAS-significant loci in specific complex diseases. We tested for enrichment of DE genes for cis-SNPs associated with diseases and phenotypes in the GWAS catalogue^47^. As many traits had a small number of associations, we first tested for enrichment within groups of similar traits, as defined in the GWAS catalogue (Figure 3, Table S6).

**Figure 3.**
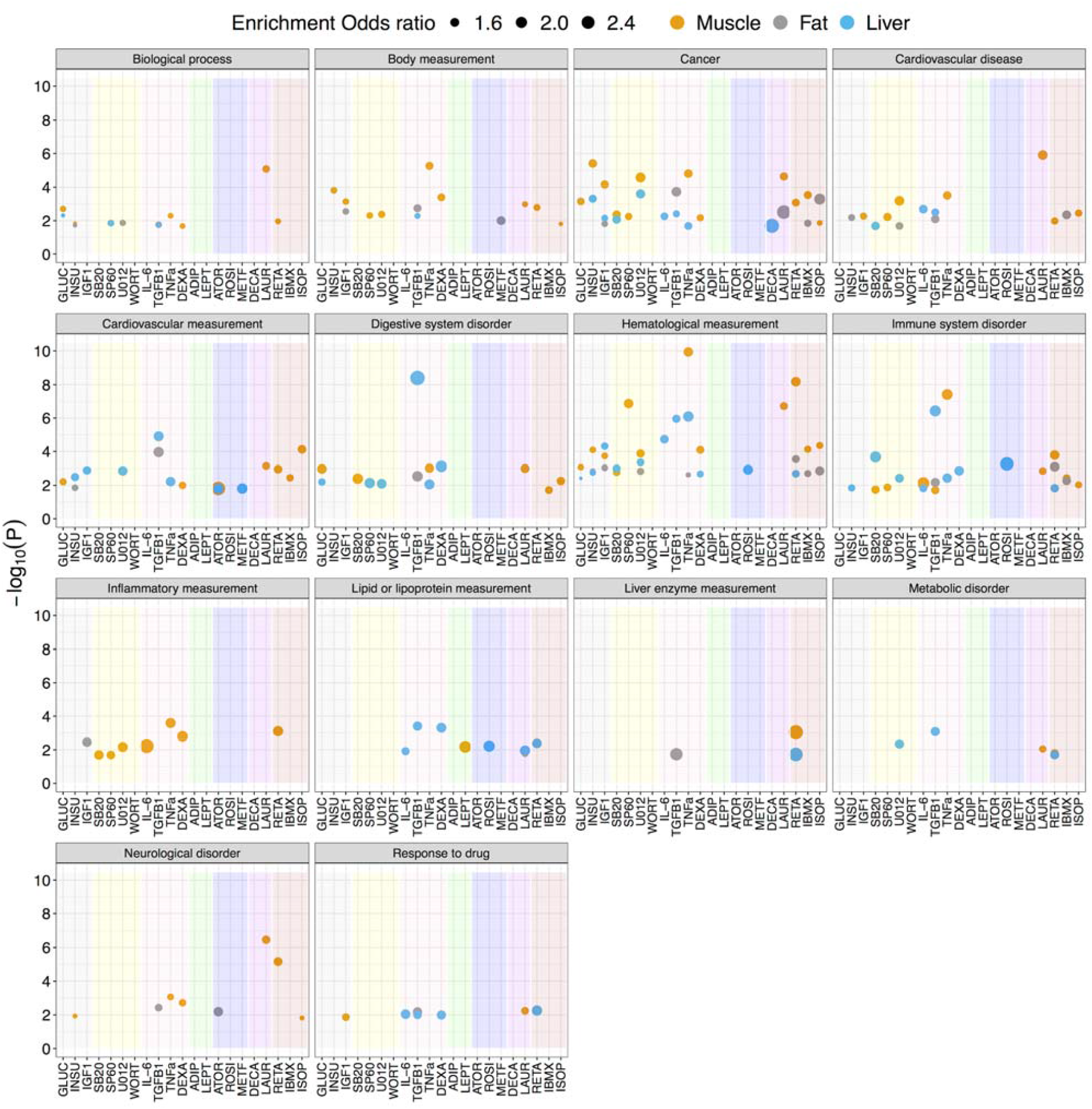
Identifying environmental perturbations impacting significant GWAS loci. GWAS enrichment results for each group of complex traits from the GWAS catalogue. Each point represents a perturbation-cell-line combination that passes the FDR<10% cut off; color of the point indicates the cell line, and the shading color within each panel indicates the perturbation category from Figure 1A. The y-axis represents the -log_10_(P-value) of the Fisher’s exact test and the size indicates the odds ratio for enrichment of GWAS hits of each group of traits from the GWAS catalogue. Numerical results are reported in Table S6. Results for specific traits, rather than groups of traits, are displayed in Figure S5.

We observed a significant enrichment for at least one perturbation and cell line across all 14 groups of complex diseases and traits tested (FDR<10%). For example, genes responsive to the effect of rosiglitazone, an insulin sensitizer known to affect plasma lipid levels^48^, were enriched within GWAS significant hits for lipid or lipoprotein measurements (OR=2.00 and P=5.98 × 10^−3^). In addition, genes responsive to the effect of retinoic acid, a metabolite of vitamin A that is synthesized in the liver and whose signaling dysregulation contributes to hepatic disease^49^, were enriched within GWAS hits for liver enzyme measurements (OR_Muscle_= 2.71 and P_Muscle_= 9.06 × 10^− 4^; OR_Liver_=2.60 and P_Liver_= 1.8 × 10^−2^). Moreover, atorvastatin and metformin, two perturbations with highly correlated DE signals (Figure 1D) known to reduce cardiovascular morbidity^50–54^, were both enriched within GWAS hits for cardiovascular measurements (OR_ATOR-Liver_= 1.75 and P_ATOR-Liver_= 1.58 × 10^−2^; OR_ATOR-Muscle_= 2.53 and P_ATOR-Muscle_= 1.52 × 10^−2^; OR_METF-Liver_= 1.86 and P_METF-Liver_= 1.56 × 10^−2^). In line with the LDSC regression-based enrichment for Crohn’s disease, we observed that genes responsive to TGF-β1 were enriched within GWAS significant hits for digestive system disorders (OR = 2.7 and P= 3.96 × 10^−9^).

More generally, we observed that GWAS hits for immune system disorders or inflammatory measurements were enriched in genes responsive to the effect of inflammatory perturbations, e.g., OR_TNFa_= 1.92 and 1.77 and P_TNFa_ = 3.84 × 10^−8^ and 2.54 × 10^−8^, for immune system disorders and inflammatory measurements respectively. Neurological disorders were also enriched for inflammatory perturbations, though to a lesser extent (e.g., OR_TNFa_= 1.35 and P_TNFa_ = 8.60 × 10^−4^). Associations with lipid or lipoprotein measurements and drug metabolism traits were enriched in genes responsive to several perturbations via liver, where most drug metabolism occurs^55^, and associations with body measurements were enriched via muscle.

For traits with a large number of GWAS hits, i.e., traits with at least 100 reported associated loci, we tested enrichments directly (Figure S5). In 14 of the 152 complex diseases and traits tested, we observed a significant enrichment for at least one perturbation and cell line (FDR<10%). For example, genes in muscle cells that were responsive to the effect of isoprenaline, a beta-adrenergic agonist with effects on cardiac muscle^56^, were enriched within GWAS significant hits for cardiovascular disease (OR=2.24 and P=2.56 × 10^−4^). In addition, consistent with the LDSCreg-based enrichment of dexamethasone for HDL heritability in liver, genes responsive to dexamethasone in liver were enriched within GWAS significant associations for total cholesterol levels (OR=3.08 and P=1.94 × 10^−4^). Lastly, genes responsive to IGF1 in the liver were enriched within significant associations for birth weight (OR=3.86 and P=5.48 × 10^−5^), consistent with prior observations of negative correlation between IGF1 levels and birth weight^57,58^.

### Environmental perturbations harbor causal genes and help inform their functions

A major challenge with GWAS data in isolation is identifying causal disease genes. Here, we assessed whether combining GWAS with relevant environmental perturbations helped to identify or reinforce causal disease genes and to inform on their molecular functions. As many of our perturbations were related to cardiometabolic traits including IR, obesity, and T2D, we tested if genes affected by our panel of perturbations harbored candidate causal genes underlying loci for seven cardiometabolic traits. To assess this, we integrated our perturbation data with results from genomic colocalization analyses of GWAS loci for these seven traits and GTEx eQTLs in visceral and subcutaneous adipose, skeletal muscle, and liver tissues^5^. We observed that genes with a transcriptional response to at least one of our environmental perturbations are enriched among the candidate causal genes, i.e., genes with high posterior colocalization probability (CLPP), for cardiometabolic traits (Figure 4A; OR=1.40, Fisher’s exact test P-value= 5.33 × 10^−4^). We next assessed whether DE genes for specific perturbation-cell line combinations were more likely to be causal, compared with non-DE genes (Figure 4B). Genes responsive to isoprenaline, SP600125 (a c-Jun N-terminal kinase inhibitor that plays an essential role in TLR mediated inflammatory responses), and TNFα in fat had significantly higher median CLPPs (FDR<10%), compared to non-DE genes (Wilcoxon test; P_ISOP_= 8.7 × 10^−4^, P_SP60_= 6.0 × 10^−3^, and P_TNFα_ = 6.0 × 10^−3^).

**Figure 4.**
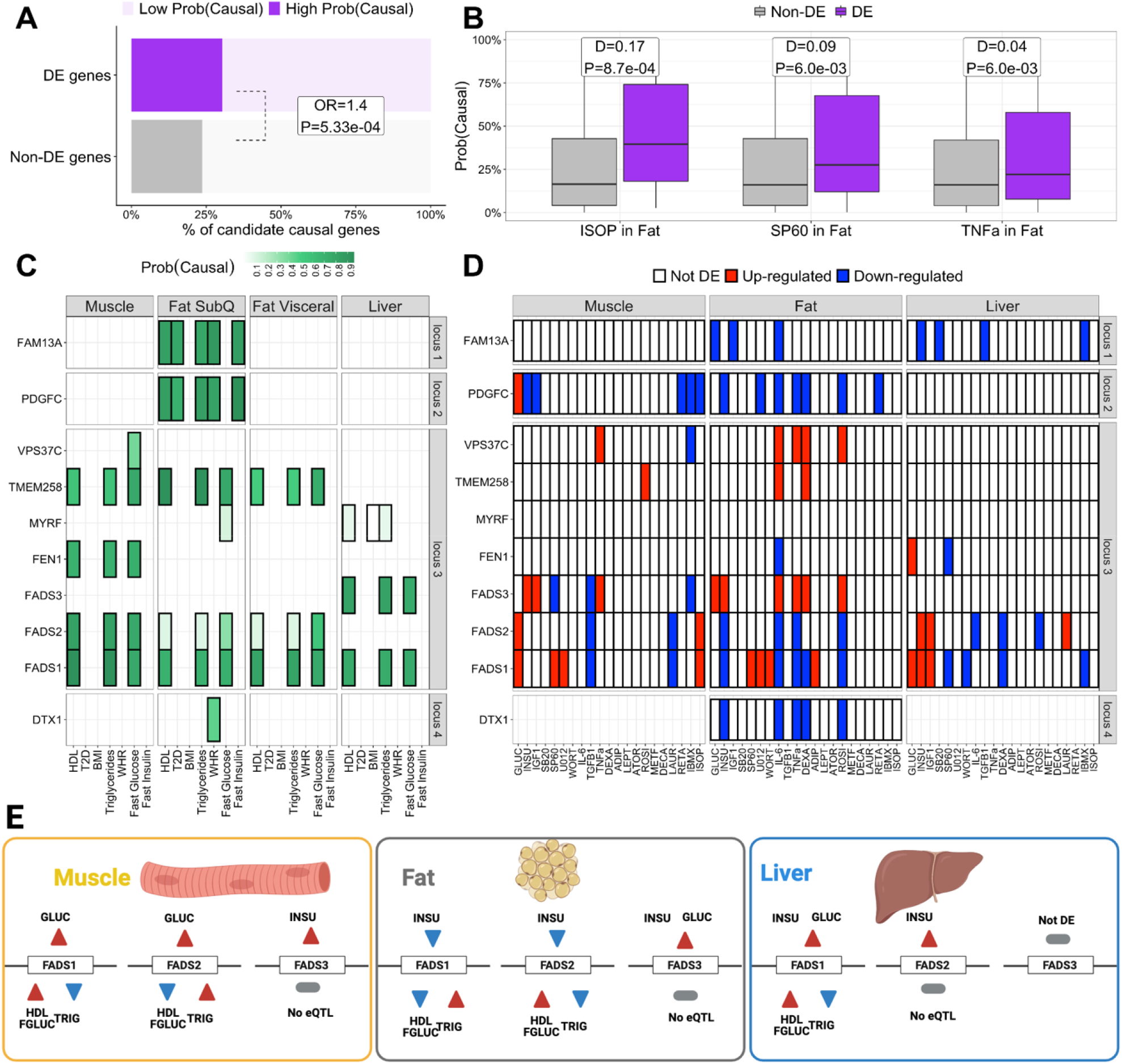
Environmental perturbations can help inform functionality of causal genes underlying cardiometabolic traits loci. **(A)** % of causal genes (High Prob(Causal)) underlying cardiometabolic traits loci that are DE (purple) or not DE (grey) in at least one perturbation and cell line. OR/P: Odds ratio and Fisher’s exact test P-value for enrichment of DE genes among causal genes, compared to non-DE genes. **(B)** Perturbation and cell line combinations with a significant (FDR<10%) difference in median (D) CLPP between DE and non-DE genes, according to the two-samples Wilcoxon rank sum test. **(C/D)** Examples of loci for which intersecting the effects of perturbations (C) with the colocalization results (D) helps inform functionality of candidate causal genes. Color indicates CLPP (C) or DE direction (D). White boxes with crosses indicate that the gene was not tested for colocalization or DE. **(E)** Effect of glucose and insulin in the expression of the three FADS genes and the effect of the expression of these genes on HDL, fasting glucose (FGLUC), and triglycerides (TRIG), the three traits for which FADS genes colocalize. The color of the triangles indicates either the effect of the perturbation on the gene (red=up-regulation, blue=down-regulation) and the effect that up-/ down-regulation of the gene has on the phenotype (red = increased phenotype, blue = decreased phenotype). DE: differential expression. CLPP: colocalization posterior probability. FDR: False discovery rate.

To explore how perturbation experiments can inform the function of candidate causal genes underlying cardiometabolic loci, we intersected the DE patterns in each cell line and perturbation with the colocalization patterns in the matched tissue. We illustrate four such examples (Figure 4C-E): three loci in which a single gene showed high CLPP and one locus with more complex colocalization patterns, with five out of seven genes in the locus showing high CLPP.

Results from the colocalization analysis associated *FAM13A* genetic variants in subcutaneous fat with several traits of interest, *i*.*e*., HDL, T2D, triglycerides, WHR, and fasting insulin (Figure 4C, locus 1). We recently described the role of *FAM13A* in adipocyte differentiation and the contribution to body fat distribution^59^. The DE patterns of *FAM13A* in our perturbation experiment (Figure 4D, locus 1) not only reinforce the role of *FAM13A* in adipose tissue but also suggest an additional metabolic function in the liver not captured by the colocalization results. The role of *FAM13A* in regulation of hepatic glucose and lipid metabolism was recently confirmed by Lin et al^60^. Another candidate gene, *PDGFC*, shows an identical colocalization pattern to *FAM13A* (Figure 4C, locus 2) and the perturbation data also supports its importance in the adipose tissue (Figure 4D, locus 2). However, the perturbation data identifies an additional role of *PDGFC* in skeletal muscle, in contrast with the role of *FAM13A* in the liver.

Another complementary example is illustrated by the colocalization for *DTX1*, which is specifically associated with WHR and subcutaneous fat (Figure 4C, locus 4), and whose expression is regulated by insulin, IL-6, TNF-a, dexamethasone and rosiglitazone in mature human adipocytes (Figure 4D, locus 4).

Finally, genetic variants in the FADS locus have been associated with HDL cholesterol, triglyceride levels, fasting glucose, and T2D^61–63^ and our colocalization analysis was consistent with these observations (Figure 4C, locus 3). However, the high amount of linkage disequilibrium, the gene density and the pleiotropy of FADS genes have challenged the dissection of individual gene effects. Particularly informative is the case of *FADS1, FADS2*, and *FADS3*, for which the DE patterns for glucose and insulin (Figure 4D, locus 3) point, among others, to a fine-tuned cell line- and perturbation-specific regulation of the *FADS* locus in the context of metabolic homeostasis (Figure 4E).

Together these results highlight the importance of perturbation experiments to contextualize GWAS associations and results from genomic colocalization analyses.

## Discussion

We have profiled transcriptional responses to multiple environmental perturbations to identify disease-relevant perturbations modulating genetic risk for complex traits and to inform functionality of causal genes. By combining gene expression data with GWAS summary statistics of complex traits, we show that heritability of multiple complex traits is enriched in regions surrounding genes responsive to particular sets of perturbation-cell line combinations. We confirmed several well-established associations, e.g., the role of TGF-β1 on asthma, and provided additional evidence for recent and less well-understood associations, e.g., the role of leptin on major depressive disorder. In addition, beyond the broad polygenic impacts of the tested perturbations, we were able to prioritize the subset of perturbations that are enriched for their impact on GWAS-significant loci in specific groups of complex diseases. We observed that environmentally responsive genes are enriched for cis-SNPs associated with a broad spectrum of diseases and phenotypes from the GWAS catalogue. Further, by integrating gene expression data with information from genomic colocalization studies, we showed that environmentally responsive genes are enriched for candidate causal genes for cardiometabolic traits, and that the effects of these perturbations in the relevant tissues further suggest context-specific molecular mechanisms of GWAS hits for cardiometabolic traits.

Our approach interrogated multiple cell lines and perturbations, but comparable applications will be limited by the cell lines and environmental perturbations that are selected, the concentrations of these perturbations, and the time length for which the cells have been exposed. Further, the use of cell lines provided the opportunity for conducting well-controlled perturbation experiments; however, it is unknown the degree to which all findings would generalize to a primary cell. For some of the diseases we considered, the studied cell lines might not represent the cell type or tissue through which disease is manifested. However, because we observed that cell lines can share transcriptional responses (Figure 1D), our study design has shown that we can identify important perturbations without the causal cell type being examined.

In conclusion, we demonstrate the advantages of large-scale characterization of transcriptional changes in diversely stimulated and pathologically relevant cells to identify disease-relevant perturbations that modulate genetic risk for complex traits. We also provide a broad resource of the dynamic transcriptional landscape in metabolic tissues. To our knowledge, this is the largest and most complete study of transcriptional effects of metabolically relevant perturbations in human fat, liver and skeletal muscle cell lines. In addition, we show that integrating GWAS and eQTL results with perturbation experiments can inform the function of candidate causal genes and prioritize genes and environmental stimuli for follow-up experiments. Combined, this work demonstrates how integrating differential expression, eQTL, and GWAS data can inform genetic and environmental components of complex disease mechanisms.

## Online Methods

### Cell culturing and perturbations

Experiments were conducted using human skeletal muscle^64,65^ (HMCL-7304), fat^66^ (Simpson-Golabi-Behmel syndrome -SGBS), and liver^67^ (HepG2) cell lines. Details on cell culturing are provided in the Supplemental Methods. Briefly, each cell line was starved for 6 hours and then treated for 2 hours with one the 21 perturbations shown in Table S1 in triplicate for each cell line -perturbation combination. We selected a stimulation window of 2 hours to allow enough time for transcriptional changes to occur, and at the same time, to minimize potential secondary responses that are not direct transcriptional effects of the selected perturbations, as previously assayed for insulin in liver and skeletal muscle cells from mice^68^. In addition, we selected the concentrations of use, shown in Table S1, based on a consensus research of available literature, particularly in the cells of interest. Last, we prepared triplicate control samples for glucose (no glucose medium) and four sets of triplicate control samples (no stimulation) for all other perturbations in each cell line, resulting in a sample size of 234.

### RNA isolation and sequencing

After stimulation, the cells were washed with PBS and collected in PureLink RNA extraction lysis buffer supplemented with 1% 2-mercaptoethanol, flash frozen in dry ice and stored at -80°C. RNA extraction was performed with PureLink RNA Mini kit (Thermo). 260/280 and RIN values were assessed prior to sequencing for sample purity and integrity. Library preparation was performed at Novogene company. Liver samples were sequenced in HiSeq 4000 (Illumina) and fat and muscle were sequenced in Novaseq 6000 (Illumina) at 150bp paired end reads.

### RNA-seq quality control

For all subsequent analyses, we focused only on expressed genes, i.e., genes which have median expression counts above 10 in at least one of the conditions (perturbation or control) within each cell line, i.e., 17,660, 17,140, and 16,722 genes for fat, muscle, and liver cells, respectively. As a measure of quality control, we looked at several RNA-Seq technical metrics (See Supplemental Methods and Figures S1-S3), e.g., RNA integrity number, % GC content, % of uniquely mapped reads etc. One sample (TGF-β1 in Fat) was dropped due to failing these quality control metrics. All remaining samples had values within proper range (Figure S1). We used principal component analysis (PCA) to identify gene expression outliers. After removing the low-quality sample mentioned above, no outliers are present based on the first two principal components (Figure S3).

### Identifying major components of variability in RNA-Seq data

We identified major components of variability in RNA-Seq data using the linear mixed models implemented in the R package variancePartition^69^ (Figure S2). We correct all subsequent analyses for all variables that explain, on average, more than 1% of expression variability in either cell line, i.e., % GC content, % exon overlapping reads, RNA concentration, % of reads marked as PCR duplicates, RNA 260/280 ratio, RIN, and % Uniquely mapped reads. For analyses done in liver cells, we also correct for sequencing batch.

### Differential expression analyses

We characterize transcriptional responses to each perturbation in each cell line using the negative binomial models implemented in the R package DESeq2^70^, adjusting for major technical components of expression variability identified in the last section. To account for multiple testing across cell lines, perturbations, and genes, we use the hierarchical error control strategies implemented in the R package TreeBH^71^ with cell line, genes, and treatments in level 1, 2, and 3, respectively. This hierarchical procedure adjusts for all the tests performed and allows us to make statements about differential expression at the gene, gene-perturbation, and gene-perturbation-cell-line level. We call a gene perturbation-specific within a cell line, if the gene is DE in that specific perturbation but not in any other perturbation in that cell line (FDR<5% at each level). A gene is assumed perturbation-and-cell-line-specific if the gene is DE in that specific perturbation but not in any other perturbation in that or the other cell lines or that specific perturbation in the other cell lines (FDR<5% at each level).

### Correlation and hierarchical clustering of transcriptional response to perturbations

We computed correlation and performed hierarchical cluster analysis of the transcriptional response to perturbations in each cell line using the test statistics from the DE analyses for all genes that were significant (FDR<5%) in at least one perturbation and cell line. The R package corrplot^72^ was used to get a graphical display of the correlation matrix and hierarchical clustering of the perturbations.

### Enrichment analyses for biological pathways

We performed over-representation analysis^73^ using the R package clusterProfiler^74^ and pathways from the *ConsensusPathDB* database^29^. We adjust for multiple testing within each perturbation and cell line using the Benjamini-Hochberg^75^ procedure.

### LD score regression analysis

We downloaded the baseline model LD scores, regression weights, and allele frequencies from https://github.com/bulik/ldsc. Annotations for each perturbation and cell line were built using the pipeline described on the LD score regression wiki and according to ^8^. Specifically, for each of the 63 combinations of 21 perturbations and three cell lines, we add 100kb windows on either side of the transcribed region of each DE gene in that combination to construct a genome annotation corresponding to that perturbation -cell line combination. Due to its unusual genetic architecture and LD pattern, we excluded the HLA region from all analyses. Z-scores for the significance of the estimated total heritability for each trait were computed as h^2^/se(h^2^), where h^2^ and se(h^2^) are the SNP-based heritability estimated and standard errors from LD score regression. Z-scores and p-values for the significance of the partitioned and conditional heritability for each trait-perturbation-cell type combination were obtained using the option --h2-cts flag. We adjust for multiple testing within each trait and cell line using the Benjamini-Hochberg procedure.

### Enrichment for diseases and traits in the GWAS catalogue

We downloaded the entire GWAS Catalogue (v1.0.2) with added ontology annotations the file showing all GWAS Catalog reported trait to EFO mappings, including the parent category each trait is mapped to on the diagram from https://www.ebi.ac.uk/gwas/docs/file-downloads. We assume the MAPPED GENE, i.e., the gene mapped to the strongest SNP as reported in the GWAS catalogue, is the GWAS gene. For the enrichment of groups of GWAS traits, i.e., EFO parent terms, we only keep the unique GWAS genes reported across all traits within each EFO parent terms. We excluded all results annotated with “Other measurement”, “Other disease”, and “Other trait” EFO parent terms as well as duplicated entries. For the enrichment of specific traits, we only test traits with at least 100 reported associated genes. To test for the significance of the enrichment, we used the Fisher’s exact test. For each perturbation and cell line combination, we use an equal number of non-DE genes matched for length and median gene expression using the R package optmatch^76^. We adjust for multiple testing across all (parent) traits, cell lines, and perturbations using the Benjamini-Hochberg procedure.

### Colocalization analysis of GWAS and eQTL effects and combination with DE signal

We performed colocalization analysis using our custom integration of the FINEMAP^77^ and eCAVIAR^11^ methods. For each GWAS and eQTL overlap (GWAS and eQTL P < 5e-8 for at least one SNP in each), we narrowed our summary statistics to the set of SNPs tested for association with both the given GWAS trait and the given QTL trait, and removed all sites containing less than 10 SNPs after this filter. Using the full 1000 Genomes dataset from phase 3 as a reference population^78^, we estimated LD between every pair of SNPs. We then ran FINEMAP independently on the GWAS and the eQTL summary stats to obtain posterior probabilities of causality for each of the remaining SNPs and combined these probabilities to compute a colocalization posterior probability (CLPP) using the formula described in the eCAVIAR method. Because the canonical CLPP score is highly conservative in regions with densely profiled, high-LD SNPs, we modified the score formula to produce an LD-modified CLPP score (Supplemental methods).

To test whether the genes DE in at least one of our perturbations and cell lines are enriched for candidate causal IR genes for at least one IR-related trait and GTEx tissue we used Fisher’s exact test. Candidate causal IR genes, denoted as High P(Causal), are defined as genes with CLPP above 40%, which corresponds to 80^th^ CLPP percentile. To test for the significance of the difference in median CLPP between DE and non-DE genes for each combination of perturbation and cell line we used the two-samples Wilcoxon rank sum test. We adjust for multiple testing using the Benjamini-Hochberg procedure.

## Author Contributions

BB, ICO, JWK, SBM, and TQ were involved in the conception and design of the work. MW provided the SGBS cells and methods for SGBS cell culture. ICO performed all the experiments. BB, MJG, and MGD performed all the statistical analyses. BB and MJG were involved in acquisition and quality control of additional data. BB drafted the manuscript with input from SBM, ICO, and JWK. All authors contributed to the interpretation of results and critically reviewed the manuscript.

## Acknowledgments

We thank Noah Zaitlen, Erik Ingelsson, Paivi Pajukanta, Eran Halperin, and Aldon J. Lusis for helpful comments and suggestions that helped to improve the quality of our manuscript.

