## Supplemental material for "An integrated approach to identify environmental modulators of genetic risk factors for complex traits"

Brunilda Balliu*^¶^

*Department of Computational Medicine, University of California Los Angeles, Los Angeles, CA, USA*

Ivan Carcamo ­Orive*^¶^

*Department of Medicine, Division of Cardiovascular Medicine, Cardiovascular Institute and Stanford Diabetes Research Center, Stanford University School of Medicine, Stanford, CA, USA*

Michael J. Gloudemans

Biomedical Informatics Training Program and Department of Pathology, Stanford University School of Medicine, Stanford, CA, USA

Daniel C. Nachun

*Department of Pathology, Stanford University School of Medicine, Stanford, CA, USA*

Matthew G. Durrant

*Department of Genetics, Stanford University School of Medicine, Stanford, CA, USA*

Steven Gazal

*Center for Genetic Epidemiology, Keck School of Medicine, University of Southern California, CA, USA*

Chong Y. Park

*Department of Medicine, Division of Cardiovascular Medicine, Stanford University School of Medicine, Stanford, CA, USA*

David A. Knowles

*New York Genome Center, New York, NY, USA*

Martin Wabitsch

*Department of Pediatrics and Adolescent Medicine, Division of Pediatric Endocrinology, Ulm University, Ulm, Germany*

Thomas Quertermous

*Department of Medicine, Division of Cardiology and Cardiovascular Institute, Stanford Diabetes Research Center, Stanford University School of Medicine, Stanford, CA, USA*

Joshua W. Knowles*

*Department of Medicine, Division of Cardiology and Cardiovascular Institute, Stanford Diabetes Research Center and Stanford Prevention Research Center, Stanford University School of Medicine, Stanford, CA, USA*

Stephen B. Montgomery*

*Department of Pathology and Department of Genetics, Stanford University School of Medicine, Stanford, CA, USA*

### * correspondence to, and

### ¶ These authors contributed equally to this work.

#### *Cell culture, differentiation and stimulation treatment*

Experiments were conducted using HepG2 (liver; ATCC) and terminally differentiated SGBS (pre-adipocytes; provided by Dr. Martin Wabitsch, Ulm University, Ulm, Germany) and HCML-7304 (myocytes; provided by Institute of Child Health, University College London) cell lines. SGBS and HCML-7304 cells were differentiated as described previously^1,2^. HepG2, terminally differentiated SGBS, and HCML-7304 cells were starved for 6 hours in EMEA with no FBS (HepG2), DMEM/F12 supplemented with pan/bio and penicillin/streptomycin but no FBS (SGBS adipocytes) or HCML growth medium (PromoCell) without supplements. For the glucose condition DMEM no glucose medium (Thermo) was used. After starvation and washing with PBS, the cells were exposed for 2 hours to the different compounds or the respective control media. The concentrations used are detailed in Table S1. All samples were generated in triplicate.

#### *RNA-seq quantification*

Reference genome (hg19) and gene model annotation files were downloaded from the UCSC genome browser website directly. Indexes of the reference genome were built using STAR^3^ and paired-end clean reads were aligned to the reference genome using STAR (v2.6.0; with default option mismatch = 10). Bam files were filtered for uniquely mapped reads, sorted, and indexed using SAMtools^4^ (v1.4.1). HTSeq^5^ (v0.11.0) was used to count the read numbers mapped to each gene (with option -m union).

#### *RNA-seq quality control*

As a measure of quality control, we looked at the following metrics: RNA integrity number, number of sequenced reads, % GC content, % of reads marked as PCR duplicates, % of exon, intron, and transcript overlapping reads, and % of uniquely mapped reads. One sample had a high % of reads marked as PCR duplicates (90%) as well as a low % of (uniquely) mapped reads (4%). This sample was excluded from further analyses. All remaining samples had RIN above 8, at least 16 million reads, an average of 51% GC content, an average of 36% of reads marked as PCR duplicates, at least 84% of their reads mapped uniquely, an average of 95%, 4%, and 98% exon, intron, and transcript overlapping reads. Moreover, for all samples, their median Spearman expression correlation (D-statistic) with other samples was at least .96 (Figure S1).

#### *Identifying major components of variability in RNA-Seq data*

To identify and adjust our analyses for major components of variability in RNA-Seq data, we computed the proportion of expression variance explained by each RNA-Seq technical metric using the R package *variancePartition* (Figure S2). Prior to computing the % of variance explained, we variance stabilized and log2-transformed the expression of each gene within cell lines using the R package DESeq2. Then, we centered and scaled each gene to have zero mean and unit variance. For each gene expressed in each cell line, we used a linear mixed model with the effect of medium, number of cells plated, plate number, sequencing batch, cell collection, differentiation, RNA extraction, starvation and treatment date as random and the effects of all other variables as fixed. Sequencing batch and number of cells plated only differed and were modelled for liver samples while differentiation date only differed and was modelled for fat and muscle samples. The number of cells plated, plate number, and cell collection, starvation, differentiation, and RNA extraction date are highly correlated with the treatment and could act as confounders of the treatment effect. To account for this, for liver cells we matched treated and untreated samples by the number of cells plated, collection and starvation date, and plate number (for most but not all treatments, see Table S1). Because of this matching, we could not correct our analyses for RNA extraction date since it was collinear with treatment status within each treated-untreated pair of samples. For fat and muscle, we matched treated and untreated samples by differentiation date and within that, by RNA extraction date, cell collection and starvation date, and plate number (for most but not all treatments, see Table S1). To adjust for all other variables, we include them in the model when testing for differential expression by treatment.

#### *Principal component analysis*

To identify gene expression outliers, we applied principal component analysis (PCA) within cell lines (Figure S3). Prior to applying PCA, we variance stabilized and log2-transformed the expression of each gene within cell lines using the R package DESeq2. Then, we centered and scaled each gene to have zero mean and unit variance. We also applied PCA to expression data corrected for all major components of expression variability, as defined in the previous section. After removing the outlier sample mentioned above and after we regress out all major components of expression variability, we do not see any outliers based on the two first principal components.

#### *LD-modified CLPP score*

The original CLPP is defined as $CLPP=\sum_{i=1}^{N} g_{i}e_{i}$, where $g_{i}$ is the probability that the *i*^th^ SNP is the causal variant for the GWAS, $e_{i}$ is the probability that the *i*^th^ SNP is the causal variant for the eQTL trait, *N* is the total number of variants at the locus. Our LD-modified CLPP score is a generalization of this score, given by ${CLPP}_{mod}=\sum_{i,j<N} g_{i}e_{j}{LD}_{ij}$, where ${LD}_{ij}$ is the LD (r^2^) between the *i*^th^ and the *j*^th^ SNP in a reference population.

**Table S1: Environmental perturbations and experimental parameters used in our experiment.** Provided as a separate excel file. The first sheet lists the perturbations used, their abbreviations and concentration, and the category in which they belong, e.g., adipokine. The second sheet lists all the experimental parameters, e.g., medium used, RNA extraction date, etc. The third sheet lists the control samples used for DE analysis for each perturbation.

**Table S2**: **Summary statistics from differential expression by perturbation analysis for each perturbation and cell line**. Provided as a separate excel file, one for each cell line, with the columns indicating the cell, perturbation, Ensembl gene ID, HUGO gene name, the average expression of the gene in the control cells (baseMean), the log2 fold change estimate (log2FoldChange, negative = down-regulated by perturbation) and standard error (lfcSE) in perturbed cells, the DESeq2 test statistic (test_stat) and p-value. The last column (significant_5prcFDR) indicates if the gene was differentially expressed (FDR<5%) in the perturbed cells in the particular perturbation and cell after adjusting for the number of tests performed across all perturbations and cell lines.

**Table S3**: **Specificity of gene expression differences across perturbations within and across each cell line**. Provided as a separate excel file, one sheet for each cell line, with the first column indicating the HUGO gene name and the next 21 columns being an indicator variable for the gene being DE (0 = not DE, 1 = DE upregulated after perturbation, and -1 = DE downregulated after perturbation) between samples treated with perturbation and untreated samples. The last column (Nr_Pert_DE) shows the number of perturbations in which a gene is DE. If Nr_Pert_DE = 1 and ADIP = +1/-1 but all other columns are zero, then the gene adiponectin-specific, i.e. it is only DE in adiponectin but no other perturbations in that cell line. The last two sheets provide DE info summary across perturbations and cells. In the first sheet (deGene_by_perturbation), each cell indicates if the gene is differentially expressed in a particular perturbation in at least one of the three cell lines (FDR < 5%). In the second sheet (deGene_by_cell), each cell indicates if the gene is differentially expressed for the cell line in at least one of the perturbations (FDR < 5%).

**Table S4**: **Summary statistics from enrichment analysis of differential expressed genes in pathways from the ConsensusPathDB-human database.** Provided as a separate excel file, one for each cell line. The columns indicate pathway name, gene ratio, BgRatio, p-value, BH-adjusted p-value, and qvalue from the over-representation analysis as well as the geneID and number of DE genes in the pathway.

**Table S5**: **Summary statistics from LDSC regression analysis**. Provided as a separate excel file; the first sheet lists a summary of the studies used and LDSCreg-based estimates of their SNP-based heritability (h^2^) and h^2^ z-score. The second sheet lists detailed LDSCreg results for each triplet of trait, cell line, and perturbation.

**Table S6**: **Summary statistics from enrichment for GWAS association analysis**. Provided as a separate excel file; the first sheet list results from the analysis of groups of related traits (Parent_Trait), as defined in the GWAS catalogue, while the second list results for specific traits. Each row corresponds to a trait or group of traits - cell line - perturbation combination. The columns indicate the specific trait (second sheet only), parent trait, cell line, and perturbation combination tested as well as the OR of enrichment with the 95% CI, the Fisher’s exact test p-value for the significance of the enrichment as well as the BH-adjusted p-value. The last four columns contain the number of genes the were “neither DE nor GWAS genes”, “DE but not GWAS genes”, “GWAS but not DE genes”, and “both GWAS and DE genes”.

**Figure S1: RNA-Seq data quality control (QC).** Boxplot of RNA integrity number (RIN), number of sequenced reads (in M), % GC content, % of reads marked as PCR duplicates, % of uniquely mapped reads, % of exon, intron, and transcript overlapping reads, and median Spearman expression correlation (D-statistic) across samples that passed QC. All samples had RIN above 8 (mean = 9.5), at least 16M reads (mean = 30M), an average of 51% GC content, an average of 36% of reads marked as PCR duplicates, at least 84% of their reads mapped uniquely (mean = 95%), an average of 95%, 4%, and 98% exon, intron, and transcript overlapping reads. For all samples, their D-statistic was at least .96. M: millions.

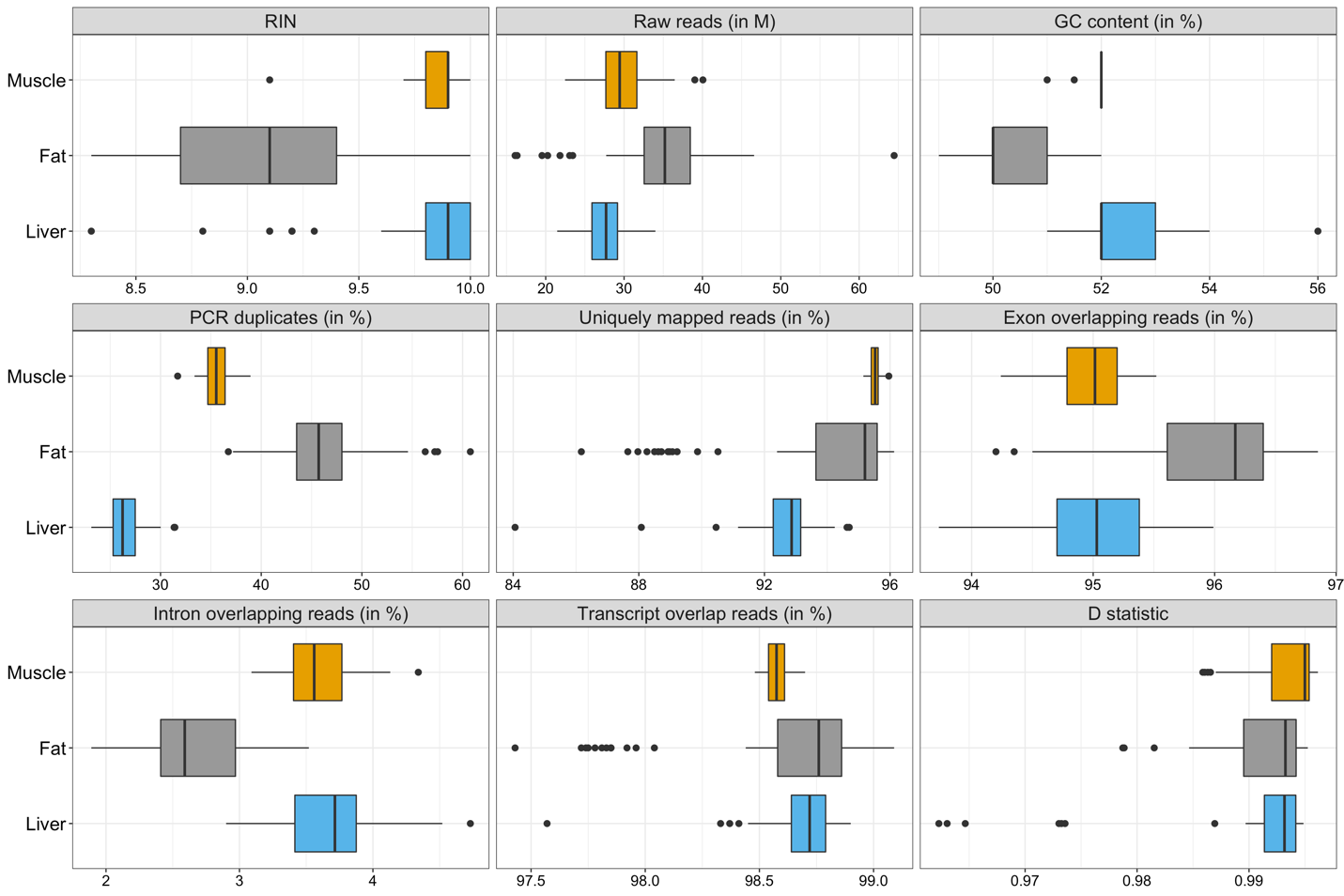

**Figure S2: Identifying major components of variability in RNA-Seq data.** Proportion of gene expression variance explained (VE) by technical metadata. Numbers next to the violin plots show the mean proportion of VE across all genes. To get the % of VE by each metadata for each gene, we used a linear mixed model with effect of medium, number of cells plated, plate number, sequencing batch, cell collection, differentiation, RNA extraction, starvation and treatment date as random and the effects of all other variables as fixed effects. Sequencing batch and number of cells plated only differed for liver samples while differentiation date only differed for muscle and fat samples.

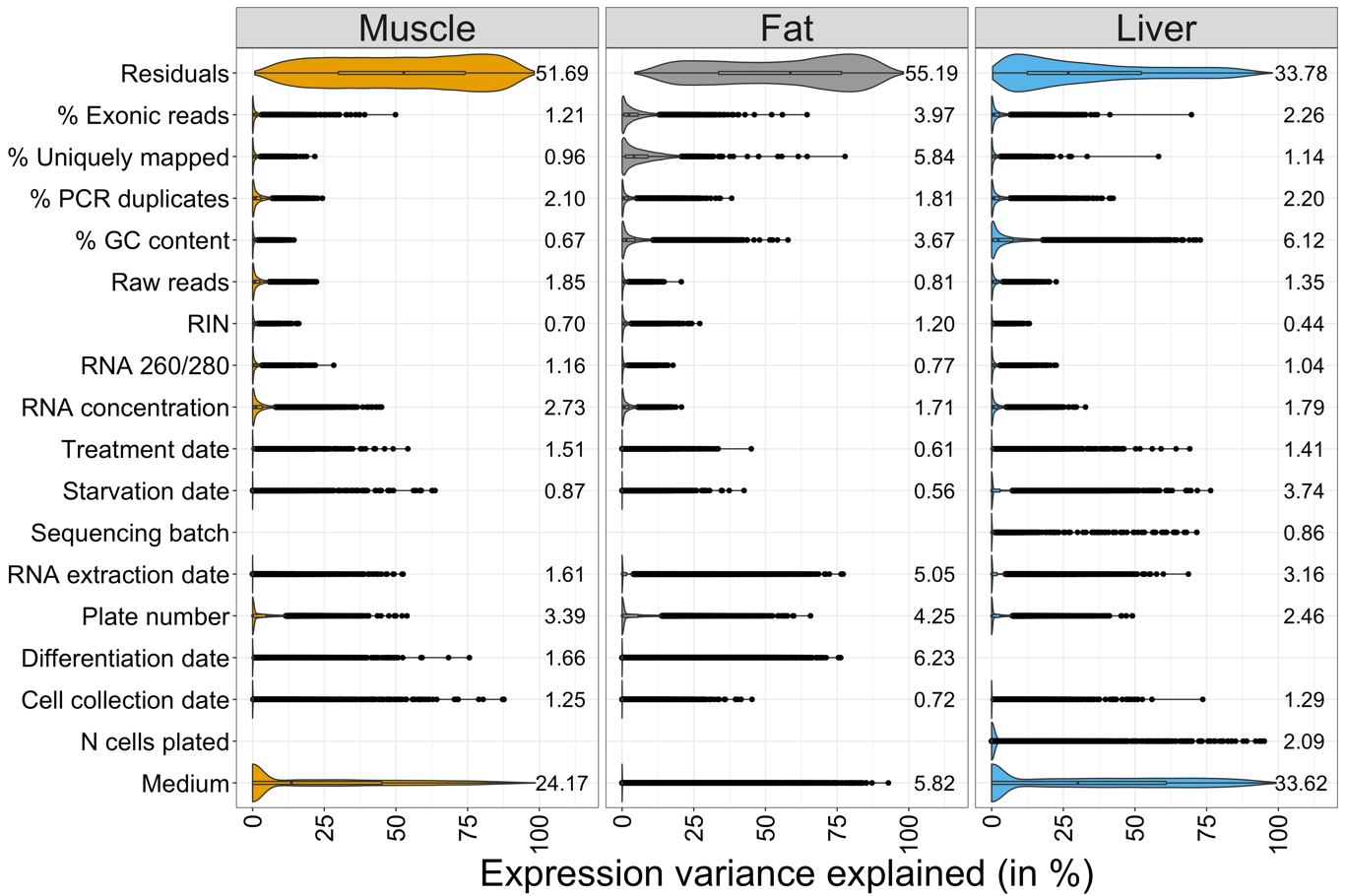

**Figure S3: Detecting outliers in RNA-Seq data using Principal Component Analysis (PCA).** Scatter plot of first two principal components (PCs). Shape indicates if the sample was glucose-related (triangle = glucose control or treated with glucose) or non-glucose related (circle). PCA was applied separately to muscle, fat, and liver samples based on raw gene expression data (top panels) and expression residuals (bottom panel) after correcting for major components of expression variability as defined in section S6 (See Figure S2).

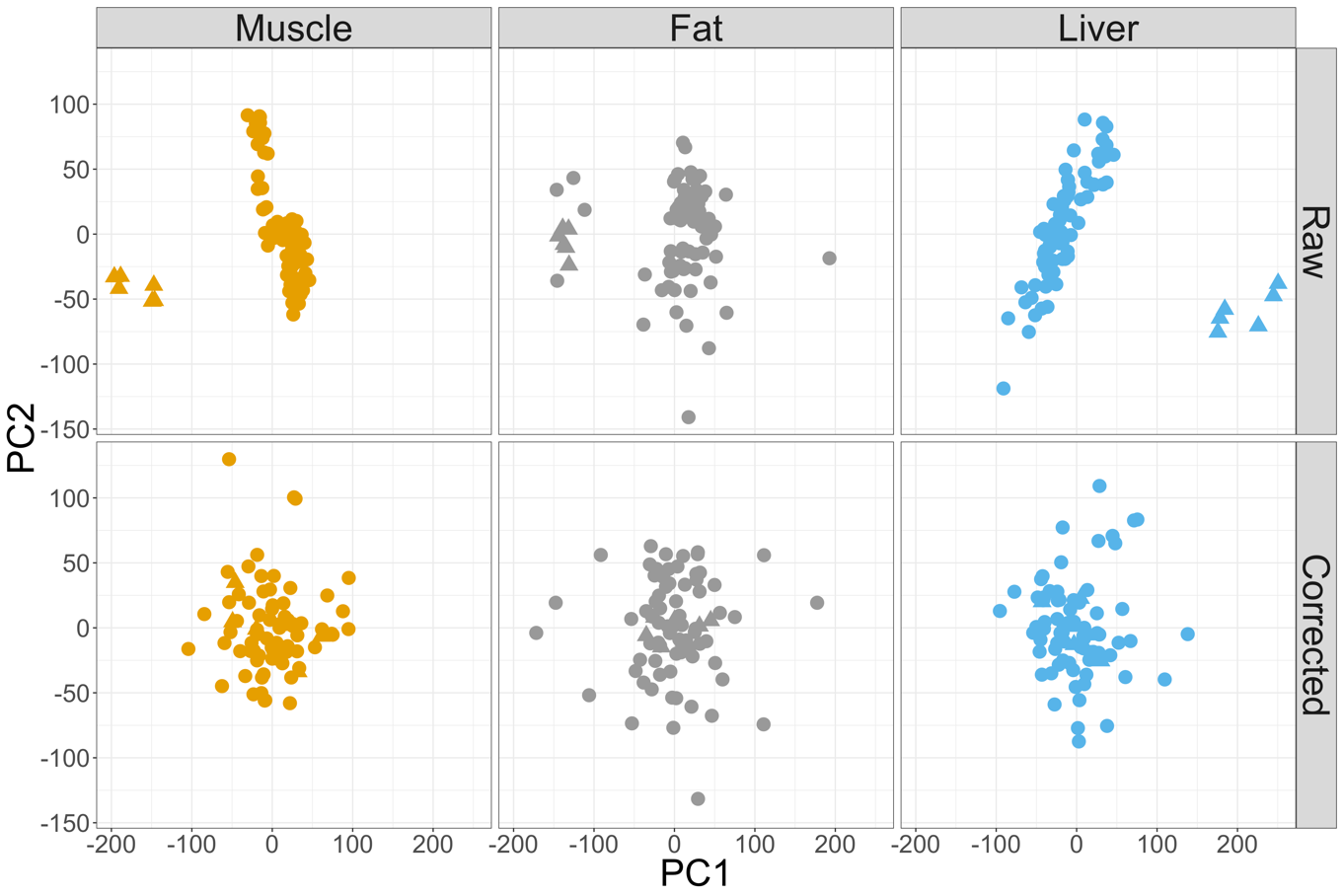

**Figure S4: Transcriptome map of 21 perturbations across human skeletal muscle, fat and liver cell line. (A)** Sharing and specificity of transcriptional responses to environmental perturbations in muscle, fat, and liver. Percent of genes that show perturbation-and-cell-line specific expression, i.e., they are DE in a specific perturbation and cell line (FDR<5%). **(B)** Pathway enrichment analysis of DE genes. The dot size represents significance of enrichment. The color represents direction of transcriptional regulation of genes in each pathway (blue: downregulated, red: upregulated). Median log_2_(FC) has been censored at (-1,1) for ease of visualization. FC: fold change. DE: differentially expressed.

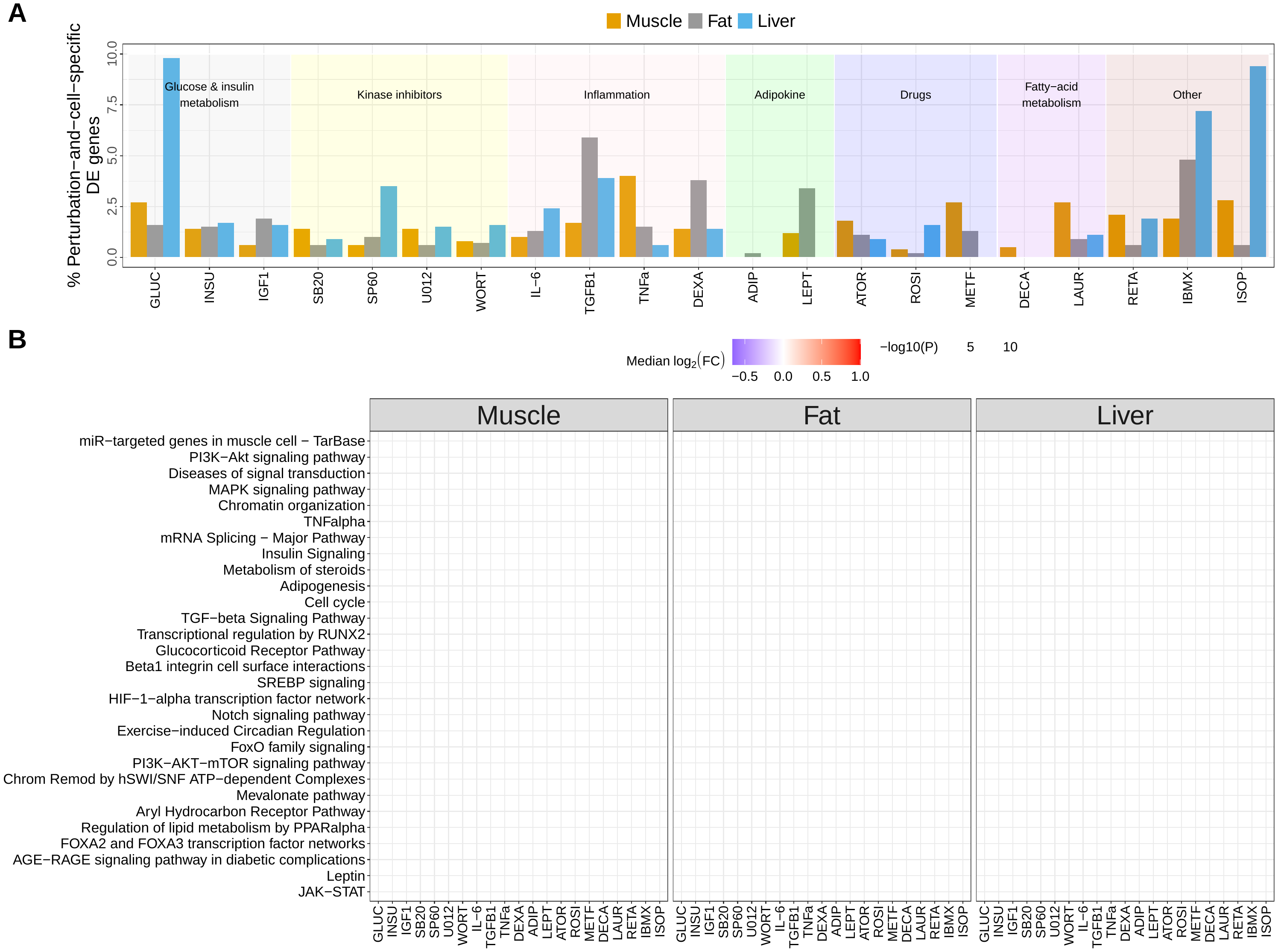

**Figure S5: Identifying environmental perturbations impacting significant GWAS loci.** GWAS enrichment results for complex traits from the GWAS catalogue. Each point represents a perturbation-cell-line combination that passes the FDR<10% cut off; color indicates the cell line. The y-axis represents the -log_10_(P-value) and the size indicates the odds ratio OR for enrichment of GWAS hits of each trait from the GWAS catalogue. The shading color within each panel indicates the perturbation category from Figure 1A. Numerical results are reported in Table SX.

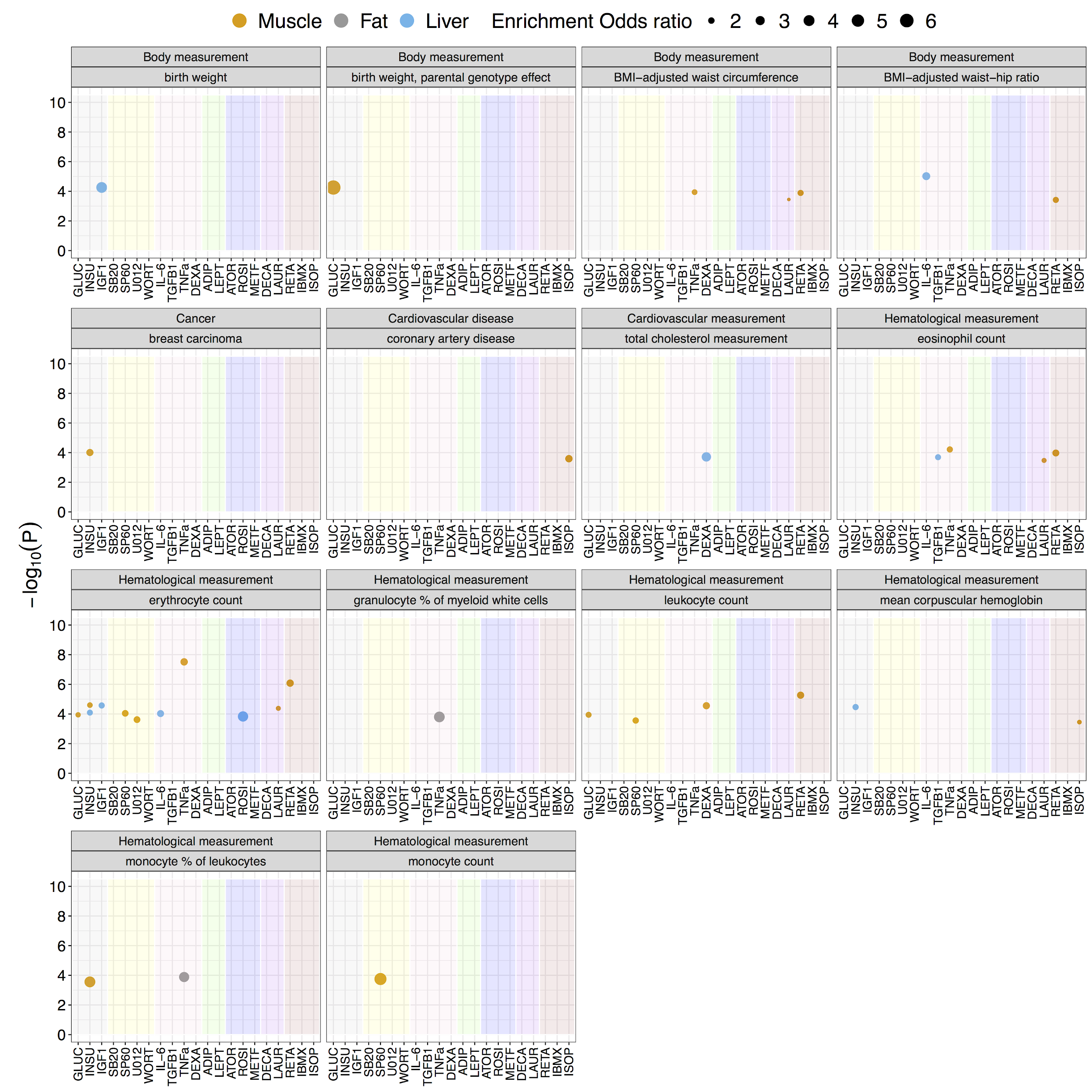
